# Fast and accurate construction of confidence intervals for heritability

**DOI:** 10.1101/031492

**Authors:** Regev Schweiger, Shachar Kaufman, Reijo Laaksonen, Marcus E Kleber, Winfried März, Eleazar Eskin, Saharon Rosset, Eran Halperin

## Abstract

Estimation of heritability is fundamental in genetic studies. In recent years, heritability estimation using linear mixed models (LMMs) has gained popularity, because these estimates can be obtained from unrelated individuals collected in genome wide association studies. Typically, heritability estimation under LMMs uses either the maximum likelihood (ML) or the restricted maximum likelihood (REML) approach. Existing methods for the construction of confidence intervals and estimators of standard errors for both ML and REML rely on asymptotic properties. However, these assumptions are often violated due to the bounded parameter space, statistical dependencies, and limited sample size, leading to biased estimates, and inflated or deflated confidence intervals. Here, we show that often the probability that the genetic component is estimated as zero is high even when the true heritability is bounded away from zero, emphasizing the need for accurate confidence intervals. We further show that the estimation of confidence intervals by state-of-the-art methods is highly inaccurate, especially when the true heritability is either relatively low or relatively high. Such biases are present, for example, in estimates of heritability of gene expression in the GTEx study, and of lipid profiles in the LURIC study. We propose a computationally efficient method, Accurate *LMM-B*ased confidence *I*ntervals (ALBI), for the estimation of the distribution of the heritability estimator, and for the construction of accurate confidence intervals. Our method can be used as an add-on to existing methods for heritability and variance components estimation, such as GCTA, FaST-LMM, GEMMA, or EMMA. ALBI is available at http://www.cs.tau.ac.il/~heran/cozygene/software/albi.html.

## Introduction

It has been known for decades that genetic variation accounts for a substantial portion of disease risk. Quantifying this portion, or estimating the heritability, had been traditionally performed using related individuals such as in twin studies [1–3]. Genome-wide association studies (GWAS) have identified thousands of genetic variants that are associated with dozens of common diseases [4, 5]. A natural approach to estimating the heritability from GWAS is to consider the total heritability explained by the identified variants. However, each of these implicated variants explains only a small fraction of the genetic component of the trait, and for many traits, even the sum total of the contributions of all identified genetic variants only explains a fraction of the heritability that was estimated from twin studies [6].

More recently, linear mixed model (LMM) approaches [7–10] have been applied to estimate the heritability from cohorts of unrelated individuals, such as in GWAS [11]. LMMs achieve considerably more accurate estimates because they can utilize all variants, and not just the variants that are statistically significant from a GWAS. Heritability estimates using LMMs are now widely utilized to understand the genetic architecture of complex traits [12]. In these studies, inferences about the genetic architecture of the trait are made by interpreting the heritability estimates.

Because there is statistical uncertainty in the estimation, a typical study uses confidence intervals (CIs) for the heritability rather than point estimates in its analysis. Unfortunately, as we show in this paper, the CIs reported by current LMM approaches are highly inaccurate (see also [13–17]). This is because these approaches make several assumptions about the data, mainly including the assumption that the estimators follow certain asymptotic distributions. In particular, estimators for traits with either significantly high or low heritability are especially biased because the estimates are near the boundaries. Additionally, these CIs may spread beyond the natural boundaries of their parameters, e.g., including negative values for heritability. As a result, these CIs are often inaccurate, difficult to interpret, or lead to erroneous conclusions.

These biases exist because estimators do not necessarily obey the conditions required for them to asymptotically follow the normal distribution. To cope with this, non-standard asymptotic theory for boundary and near-boundary maximum likelihood estimates has been developed for independent data (e.g., [18–20]), while others [21] have extended the theory to the case where the phenotype can be partitioned into sufficiently many independent vectors. Unfortunately, these conditions typically do not hold for genomic datasets. First, only a single observation is typically drawn from the distribution. Second, the parameter space is bounded, as the heritability is only defined between zero and one, and the true heritability value may lie on the boundary or near it. Therefore, an alternative approach is required.

The disagreement between the theoretical distribution of estimators and their empirical distribution results in consequences which are not limited to CIs only. One important property of the theoretical distribution of estimators, when the true parameter value is near the boundary, is a high probability for a boundary estimate [19, 20]. This is in line with a plurality of reports in the literature, in which the heritability of a phenotype is often estimated to be zero or one [14, 17, 22–24] using restricted maximum likelihood (REML) estimators, which are the state-of-the-art method in heritability estimation.

Previous approaches to remedy these issues have taken several directions. Visscher et al. [25] studied the heritability estimator in a range of pedigree- and marker-based experimental designs. While they derived an analytical expression for the variance, their method assumed that the heritability estimator follows a normal distribution. Inaccuracies in CI coverage probabilities have also been reported in longitudal studies, leading some authors to suggest exhaustive hierarchical bootstrapping schemes, e.g., [26]. Recently, Furlotte et al. [27] studied the uncertainty in heritability estimates and suggested its quantification using the Bayesian posterior distribution of the heritability value, conditioned on the observed phenotype. Other works have suggested extending the REML estimation method with Bayesian priors, e.g., [22, 24]. It has also been suggested to replace the asymptotic normality assumption with the asymptotics developed for the non-standard boundary case [28]. However, a large sample size is still required for these approximations to be effective. Finally, alternative statistics have been suggested as a basis for building CIs, such as a quadratic function of the estimator for random effects [29], or a ratio of linear combinations of quadratic functions of the phenotype [14, 30]. Unfortunately, all of these methods either assume asymptotics which typically do not hold in practice, or do not pertain to REML estimation.

In this paper, we present a novel approach for accurately building heritability CIs using LMMs. Instead of an asymptotic approximation, our method uses a parametric bootstrap test inversion approach to construct CIs via sampling phenotypes, performing heritability estimation on the sampled phenotypes, and using these estimates as a basis for CI construction [31]. As a naive implementation of this approach would be computationally prohibitive, we present a highly accurate approximation that allows us to efficiently construct the intervals. We demonstrate our approach on both the Genotype-Tissue Expression [32] and the Ludwigshafen Risk and Cardiovascular Health (LURIC) [33] datasets. An implementation of our approach, which we call Accurate LMM-Based confidence Intervals (ALBI), is available at http://www.cs.tau.ac.il/~heran/cozygene/software/albi.html.

## Results

### Heritability estimates and confidence intervals

Heritability estimates from population samples can be computed from estimates of the parameters of the linear model **y** = *μ* + **u + e**, where *μ* is the population mean, 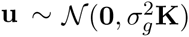 is the genetic component of the trait, and 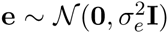 is the environmental component. **K** is a kinship matrix capturing the genetic relatedness between individuals in the sample, constructed from their genotypes. Heritability is defined as 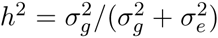. Similarly, under certain assumptions (see Methods), the heritability estimate is 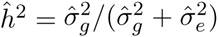. This captures the proportion of the variance of the trait that corresponds to the genetic component.

To quantify the amount of uncertainty in heritability estimates, all current methods report the standard error of their estimators. The main application of these reported standard errors is to imply the construction of CIs for the true genetic and environmental variance components 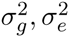, or for the true heritability, *h*^2^. These CIs are based on the assumption that the estimator is approximately normally distributed. Given an estimated value, 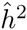, a standard error estimate, *ê*, and a required confidence level, 1 − *α* (e.g., 95%), the CI is constructed as 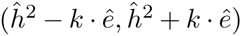, where *k* = Φ^−1^(1 − *α*/2), the (1 − *α*/2)-th quantile of the standard normal distribution. For example, a 95% confidence level CI is computed as 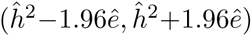. The standard error is calculated using the inverse Fisher information matrix of the estimators, or using similar approaches (for details, see e.g., [34]).

### Current methods of building confidence intervals are inaccurate

We investigated the accuracy of such CIs, using the standard error estimation performed by GCTA [35] (a popular software for heritability analysis using LMMs), which employs the approach of assuming a normal distribution for the estimator. Suppose we draw a phenotype vector from the the distribution assumed by the LMM. By the definition of a CI with a confidence level of, e.g., 95%, if we were to repeat such a draw multiple times with the same heritability value *h*^2^, and compute a CI from each draw, then these CIs should cover *h*^2^ 95% of the times. For a range of confidence levels (70%,…, 95%), we used GCTA to build a CI based on the normal approximation for that level. We then checked the percentage of times in which the CI covers *h*^2^, as a function of *h*^2^. Figure 1 uses kinship matrices from two real studies, and shows that for a wide range of *h*^2^ values, and particularly for extreme values (small or large), these normal CIs are largely inaccurate. The two studies are the Genotype-Tissue Expression (GTEx) study [32] and the Ludwigshafen Risk and Cardiovascular Health (LURIC) study [33]. The GTEx project is a sample and data resource designed to study the relationship among genetic variation, gene expression, and other molecular phenotypes in multiple human tissues. It provides a collection of multiple different tissues per donor, along with their genotypes. LURIC is targeted to contribute to the identification and assessment of environmental and genetic factors for cardiovascular diseases. Genotypes and lipid profiles are available for patients hospitalized for coronary angiography between 1997 and 2000 at a tertiary care center in Southwestern Germany (see Methods).

**Figure 1.**
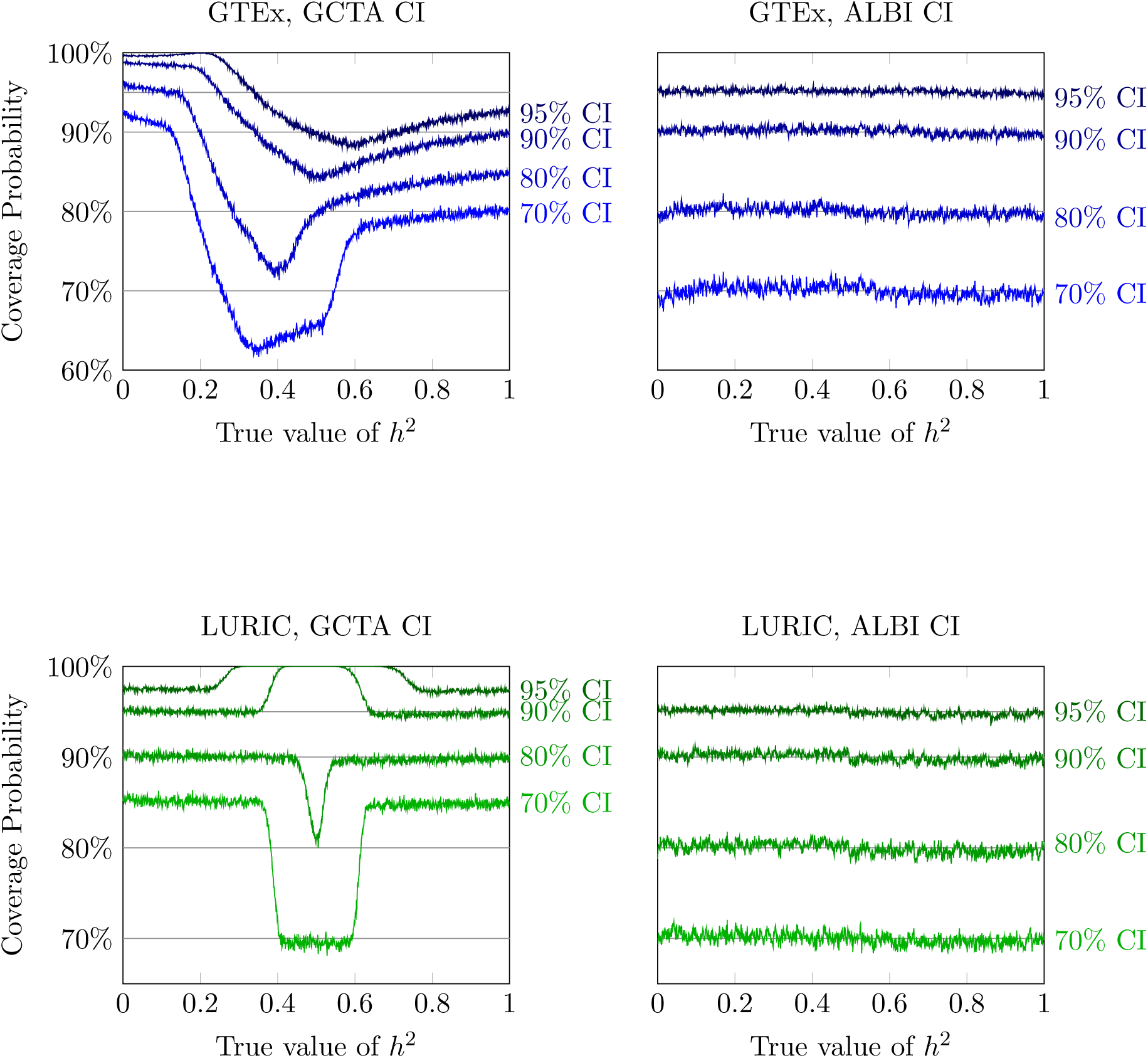
Accuracy of CIs. The left panes depict the true coverage probability of GCTA’s confidence intervals (CIs), based on the normal approximation, on the GTEx and LURIC datasets. The right panes depict the coverage probabilities of the ALBI CIs. The coverage probabilities are shown for CIs designed to have coverage probabilities of 70%, 80%, 90% and 95%. GCTA’s CIs are often far from the correct confidence level, while ALBI’s achieve accurate coverage.

The results of Figure 1 imply that the accuracy of the standard normal CIs depends both on the dataset and on the true heritability of the trait. To further demonstrate this, for each value of a true *h*^2^, we compared the empirical standard deviation of the REML estimator of 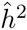 to the average of the standard error reported by GCTA. Here too, we found that these standard errors are inaccurate, especially at low and high true *h*^2^ values, with an error of a multiplicative factor of up to ×1.7 (see Figure 2). As a point of comparison, we additionally studied the Northern Finland Birth Cohort study (NFBC) dataset [36], where the extent of these biases is much more limited.

**Figure 2.**
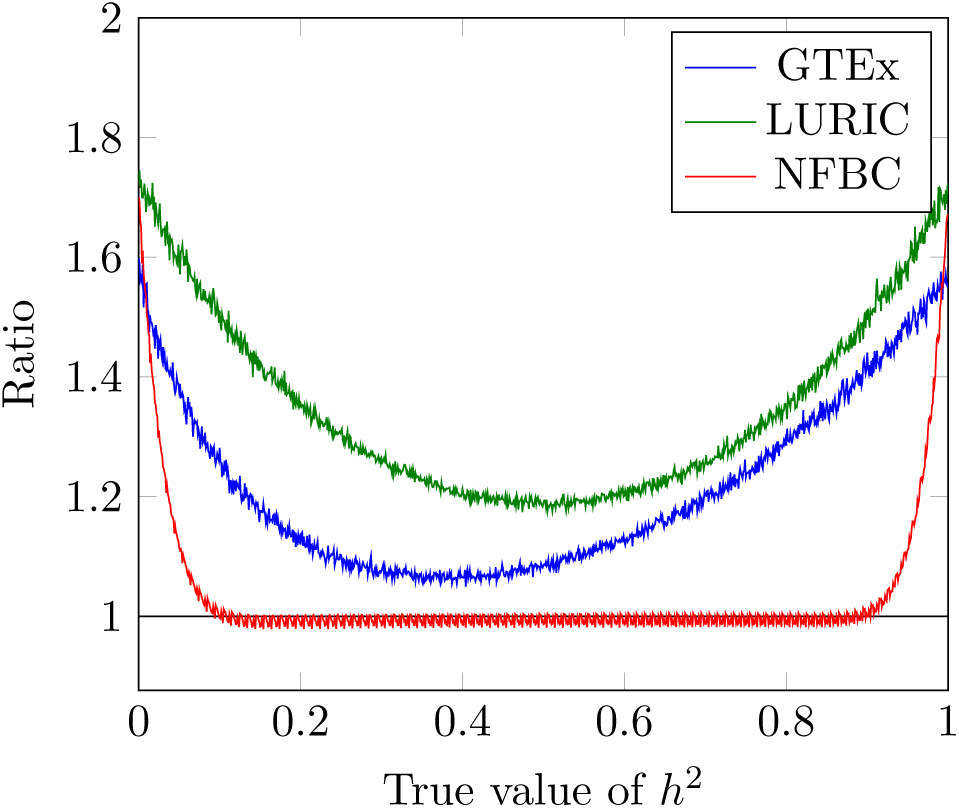
Accuracy of Standard errors. The ratio between the mean standard error derived from GCTA, and the empirical standard deviation of the REML estimator 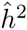, as a function of the true *h*^2^, for the studied datasets. The discrepancy is high, especially for GTEx and LURIC, with ratios up to ×1.7.

To demonstrate that the above issues are due to the normal approximation for the distribution of the estimator, we repeatedly sampled phenotypes for the studied datasets, assuming a certain heritability, and examined the distribution of the estimators. We observe that, indeed, the distribution is often far from normal, showing skewness and high probabilities for 0 and 1. Figure 3 illustrates this for several values of true *h*^2^, when 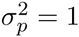, *β* = 0*_p_*. In line with the findings presented previously, the distributions from GTEx and LURIC violate the normality assumptions, while NFBC follows the normal distribution more closely.

**Figure 3.**
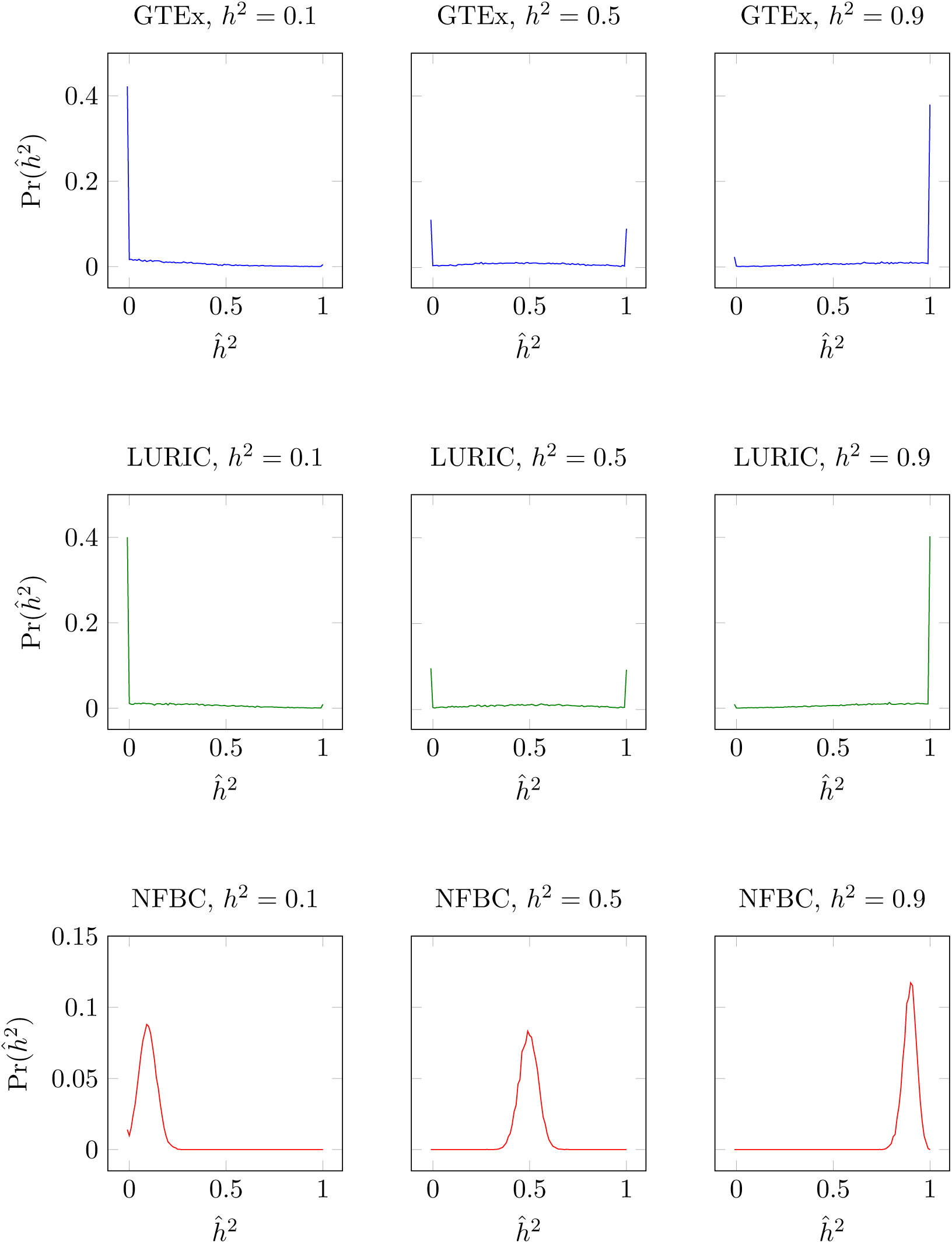
Distributions of the heritability estimator. The density of 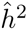 for *h*^2^ = 0.1, 0.5, 0.9 in the studied datasets, under the LMM. Since the distribution of 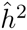 is continuous in the open interval (0, 1), we divide this range into equally sized bins of a fixed size (0.01), and instead estimate the probability mass function of a random variable taking values in the set {0, (0, 0.01], (0.01, 0.02],…, (0.09, 1), 1}. Estimator distributions are often far from being normal, hence the normal approximation seems highly questionable.

The high probability of boundary estimates 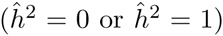, evidenced in Figure 3, is strongly reflected in published results from applied research. For example, 55% of the gene expression profiles in the GTEx data were estimated to have heritability of 0, and 15% of profiles to have heritability 1. Similarly, we calculated the heritability of expression profiles of lipids in the LURIC dataset, estimating 42% of the profiles to have 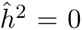 (none had 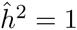). Such results are often difficult to interpret. It is not clear whether the prevalence of 0 heritability estimates suggests a true underlying zero value of heritability (*h*^2^ = 0), and therefore that a genetic variance component does not exist, or that this estimation is due to statistical noise or possibly a result of an underlying numerical error.

We can use the sampling approach described above to characterize, for a given kinship matrix, the probability that the heritability would be estimated as either 0 or 1 given the true heritability. This probability, as a function of the true heritability, is shown in Figure 4. As we can see, for the GTEx and LURIC datasets, these probabilities are high for a wide range of heritability values, which explains the large number of observed heritability estimates that equal zero. On the other hand, the NFBC dataset has a low probability of obtaining a zero estimate, unless the true heritability is small (up to ~0. 15).

**Figure 4.**
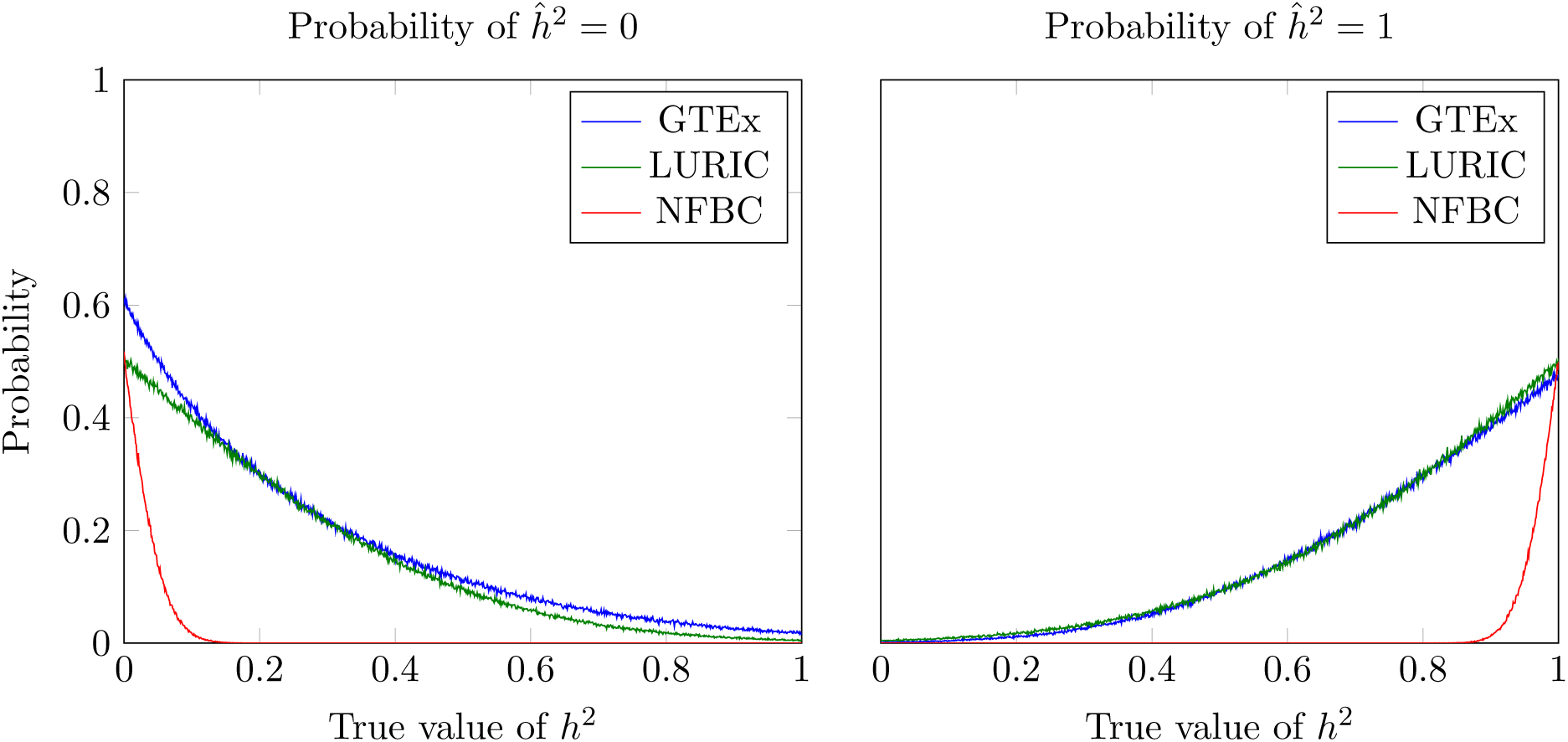
Probability of boundary heritability estimates. The probability of estimating 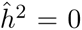 or 1, for all possible values of true underlying heritability values *h*^2^, for the studied datasets. For GTEx and LURIC, the probability of 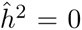 (1) is high, especially for small (large) values. For NFBC, this phenomenon is limited.

### Accurate LMM-based confidence intervals

Motivated by these inaccuracies, we have developed a novel method, which we call Accurate *LMM-B*ased confidence *I*ntervals (ALBI), for rapidly estimating the distribution of the REML heritability estimator and for computing accurate heritability confidence intervals. We give an overview of the method and its properties here. For the full description see Methods.

An explicit calculation of the estimator distribution, via the parametric bootstrap [37], can be performed as follows. For a given heritability value *h*^2^, a large number of phenotype vectors is drawn according to the LMM, and the REML estimator 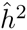 is computed for each such vector. This process can be repeated for a grid of feasible *h*^2^ values, as needed, and we can use as many bootstrap replicates as needed in order to achieve a desired accuracy. Unfortunately, this brute-force method is often a prohibitively time-consuming procedure (see Table 1).

**Table 1.**
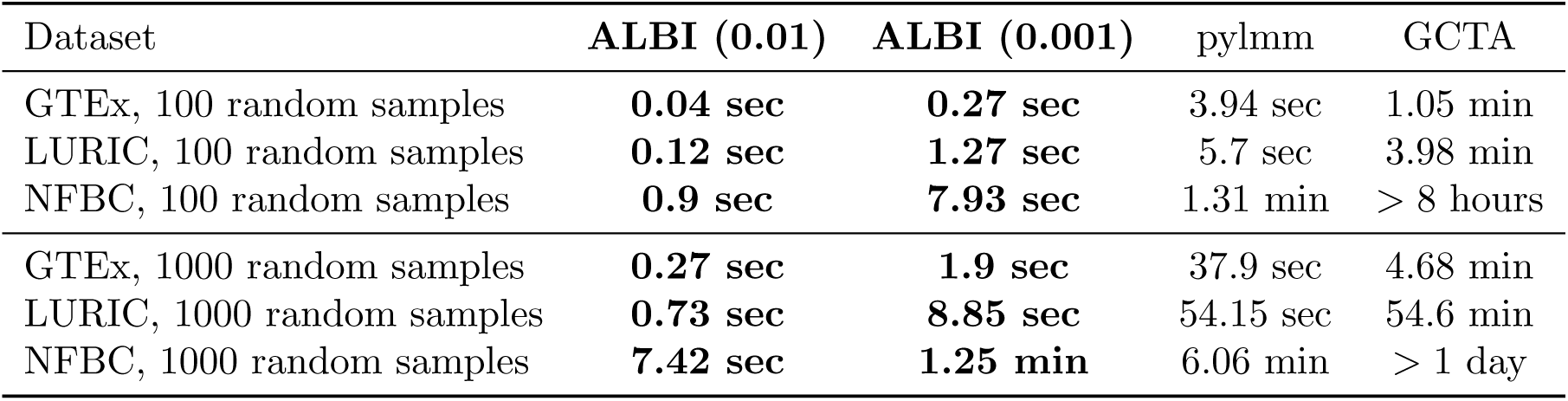
Benchmarks. Running times of ALBI vs. other brute-force methods. We compare the computational costs of ALBI to that of pylmm [13] and GCTA [1] (see Methods). We generated the estimator distribution for *h*^2^ = 0.5. For ALBI, we estimated the distributions at a precision of 0.01 and 0.001, using 100 and 1,000 random samples. For pylmm and GCTA, we generated 100 and 1,000 random phenotypes. GCTA did not converge for ~7% of the random samples. Running times are reported for the datasets of GTEx (185 individuals), LURIC (867 individuals) and NFBC (5,236 individuals).

To address this, we have developed a fast analytical approximation, which consists of the following elements. First, it depends only on the eigenvalues of the kinship matrix. Second, under several common scenarios, it gives a closed-form formula for the derivative of the restricted log-likelihood function, which can then be easily calculated. This formula also allows for the evaluation of the probability of the estimate falling within an interval without the need to estimate the entire distribution. Finally, all operations are performed using the eigenvectors of the kinship matrix, which allows for a computational complexity linear in the number of samples. Our approximation is highly efficient (Table 1) and it provides an excellent match to the brute-force estimation (see Methods).

Once we have the estimator distribution for any possible value of the true heritability, we can efficiently construct accurate CIs, based on the duality between hypothesis tests and CIs. For each true value of *h*^2^, we select a subset of 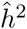 that has sampling probability 1 − *α*, according to the respective estimator distribution, and define it to be the acceptance region for that value of *h*^2^. The CI for a value 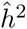 is the interval containing all values of *h*^2^ for which 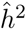 does not imply the rejection of the hypothesis of the true heritability value being *h*^2^, with a confidence level of 1 − *α*. As a comparison, we show the result of the CIs derived with ALBI (Figure 1). These CIs achieve the desired confidence level accurately.

Using ALBI, and comparing our accurate CIs to those of methods employing the normal approximation, we observed that CIs derived from the normal approximation are often too wide. This is problematic in scenarios where, for example, one is interested to know if the value *h*^2^ = 0 is included in the CI, 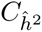. It may then be the case that 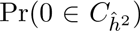 is estimated to be too high, causing the researcher to infer that a phenotype may have zero heritability, often erroneously. Indeed, 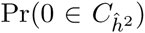 is significantly higher in GCTA than in the ALBI CIs, up to ×4.1 times in GTEx (see Figure 5).

**Figure 5.**
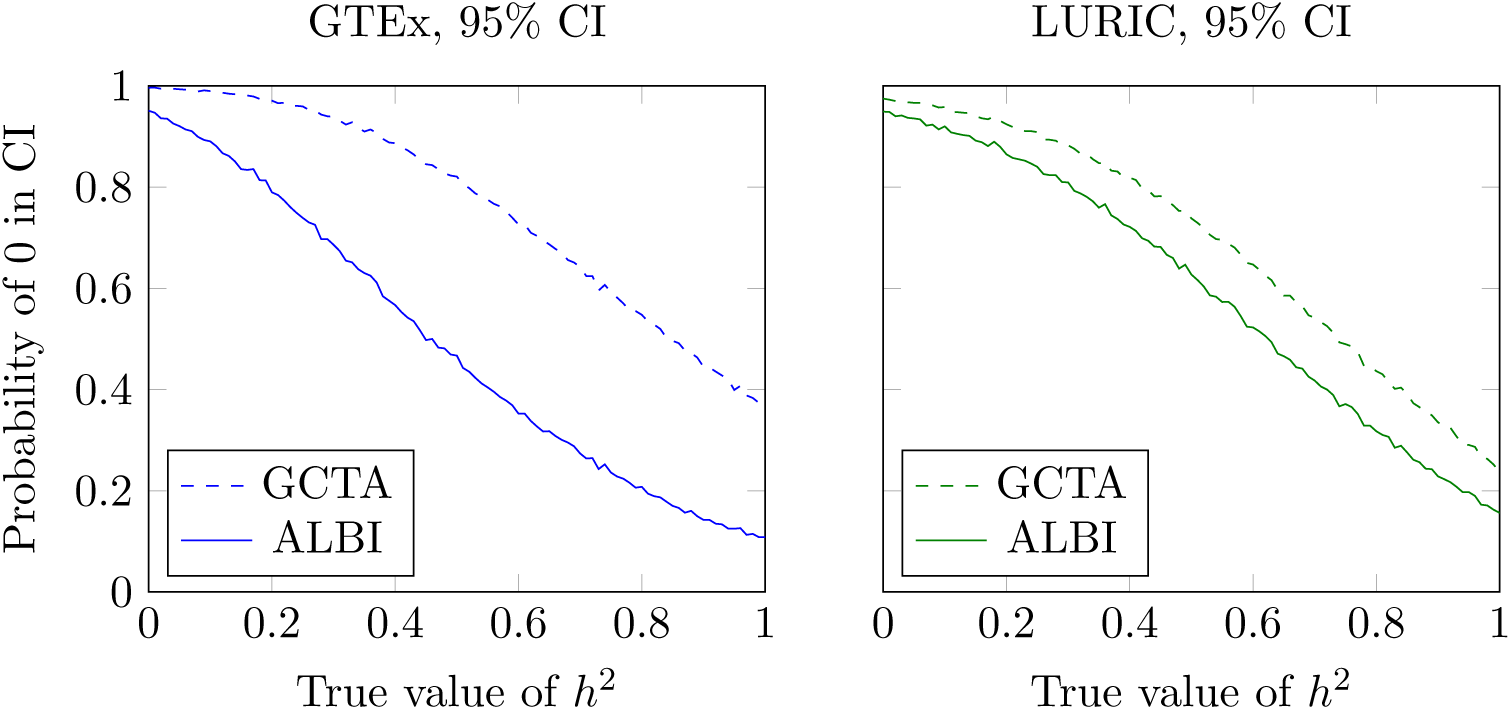
Probability of zero heritability in CI. The probability of *h*^2^ = 0 being included in the CI, as a function of the true value of *h*^2^, for the GTEx and LURIC datasets. These probabilities are shown for GCTA’s CI and ALBI’s CI, designed to have a confidence level of 95%. It can be seen that CIs derived from the normal approximation tend to include *h*^2^ = 0 more than necessary.

### A fast preliminary assessment of the accuracy of the normal approximation

As we have demonstrated, the accuracy of the normal approximation of the estimator greatly depends on the kinship matrix of the cohort used to make the estimates. We can use ALBI to efficiently compute the probability of estimating a heritability of 0 or 1, given the true heritability value, and generate the data shown in Figure 4. ALBI can compute these probabilities without the need to repeatedly sample the phenotypes and without reestimating the heritability each time, and thus it is highly efficient. Therefore, we can use these estimated probabilities to quantify how accurate we expect the normal approximation of the heritability estimates to be for a given dataset, prior to explicitly estimating the CIs for each of the estimates. For example, looking at Figure 4, it is clear that heritability estimates in the NFBC data will be much more accurate than in the GTEx or LURIC datasets.

## Methods

For clarity of presentation, we begin by defining the heritability under the linear mixed model, providing an outline of the REML estimation method, and detailing some important properties of the likelihood function. Finally, we present ALBI, our method of calculating the heritability estimator’s distribution and constructing accurate confidence intervals for heritability.

### The linear mixed model

We consider the following standard linear mixed model (see [38] for a detailed review):

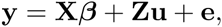

where *n* is the number of samples, y is a *n* × 1 vector of random variables, X is a *n* × *p* matrix of covariates (possibly including an intercept vector 1*_n_* as a first column), *β* is a *p* × 1 vector of fixed effects, **Z** is a *n × m* design matrix, **u** is a *m* × 1 vector of random effects, and **e** is a vector of errors. We assume **u** and **e** are statistically independent and distributed normally as 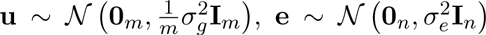. Define 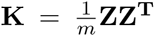. Under these conditions it follows [11] that:

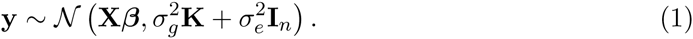

Typically, **y** is a vector of phenotype measurements for each individual, **X** is a matrix of covariates (e.g., an intercept term, sex, age) and Z is the standardized genotype matrix, i.e., columns have zero mean and unit variance. **K** is commonly called the kinship matrix, or the genetic relationship matrix, and is estimated from the genotypes as 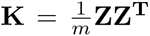 (see A.1).

The heritability is defined as the proportion of total variance explained by genetic factors [39]:

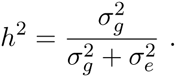

Defining 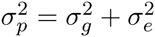, Equation (1) becomes:

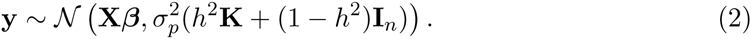

### Heritability estimation with ML and REML

The most common way of estimating *h*^2^ is REML estimation (see A.2). For completeness, we give an overview of the REML estimation method. REML consists of maximizing the likelihood function that is associated with *n − p* linearly independent contrasts [40]. The restricted log-likelihood function is, up to additive and multiplicative constants:

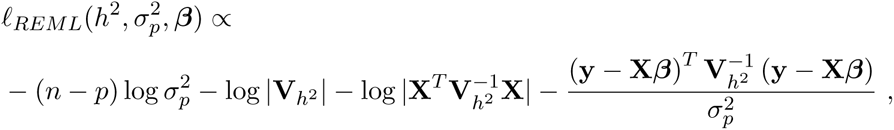

where 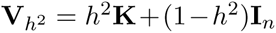. For a fixed *h*^2^, the values of 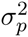 and *β* that maximize *ℓ_REML_* can be derived analytically, and plugged back in to derive a profile restricted likelihood function *f_REML_*, which is a function of *h*^2^, that is maximized instead.

In Appendix A.3, we show that the distribution of 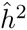 depends only on *h*^2^, and is invariant under changes to 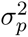 and *β*. We may therefore limit our study to the 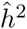 estimator alone, in the special case of fixed 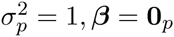, which substantially simplifies the problem.

### Estimating the distribution of REML estimators with parametric bootstrap

We now turn to describe a method to estimate the distribution of REML estimators. We begin with a direct calculation, and then describe a faster approximation.

#### Direct parametric bootstrap

For a fixed value of *h*^2^ (and assuming 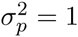, *β* = 0_p_), we can estimate the distribution of 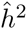 with a parametric bootstrap method. Since the distribution of 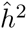 is continuous in the open interval (0,1), we divide this range into equally sized bins of a fixed size (0.01), and instead estimate the probability mass function of a random variable taking values in the set {0, (0, 0.01], (0.01, 0.02],…, (0.09, 1), 1}.

Explicitly stated, the method consists of the following steps:

1. Random sampling: Draw *N* (e.g., 10,000) phenotypes 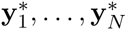 from the multivariate normal distribution 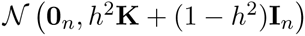.
2. REML estimation: Calculate the REML estimates 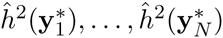 for each of these phenotypes, using a software package such as GCTA.
3. Density estimation: For each one of the bins above, count the proportion of estimates 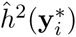 that fall in that bin; similarly, compute the fraction of estimates evaluating to a boundary estimate 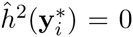 or 1. Use these fractions as an estimate of the density of 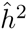 for this value of *h*^2^.

In what follows, we discuss how to perform each one of these steps more efficiently.

#### Step 1: Random sampling

Drawing a vector **y** from the distribution 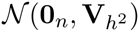 may be done by drawing a vector of standard i.i.d. normal variables 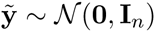, and calculating 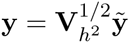. Any statement about **y** can then be restated in terms of 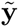, or further in terms of a vector 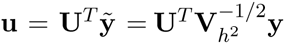, where **U** is a matrix whose columns are the eigenvectors of **K**. Note that since **U** is orthonormal, **u** is also distributed 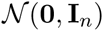. Therefore, instead of drawing multiple phenotypes 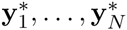, we draw u_1_,…, 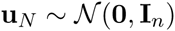, and rephrase later stages in terms of these **u**-s. Using **u** instead of 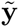 simplifies further calculations, and additionally does not require expensive matrix multiplications.

#### Step 2: REML estimation

**Local evaluation**. Instead of finding the global maximum of the restricted profile log-likelihood function *f_REML_* directly, we employ two changes: (i) we search for local maxima instead of the global maximum; (ii) we use the derivative 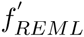 instead of *f_REML_* itself.

The main advantage of this approach is that in order to evaluate if a point estimate 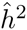 is a maximum, we need only check a local condition at this point. For example, to check if 0 is a local maximum, we simply evaluate if 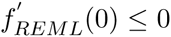, so evaluating all other points 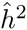 across the range is not required. In theory, it is possible that multiple local maxima exist, in which case we compare the likelihood itself directly at these points. However, in practice we have observed that this multiplicity happens only rarely, and even an arbitrary decision between local maxima does not noticably hurt CI accuracy.

As we are estimating the distribution of 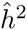 at a specified resolution, we are in fact more interested in the question of the estimate 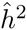 being inside an interval. Using the derivative, it is simple to check if an interval (*c*_1_, *c*_2_) contains a local maximum; If 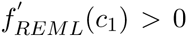 and 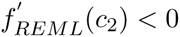, then there exists at least one 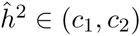 which is a local maximum. While this is only a sufficient condition for the existence of a local maximum within the interval, we find that when the interval is sufficiently small (e.g., of width 0.01), it is also a necessary condition. In addition, the probability of having multiple local maxima inside a small interval is negligible. Therefore, evaluations are only necessary at grid points.

**Closed-form formula**. From the discussion above, it follows that we are interested in the condition of evaluating the derivative of the profile restricted log-likelihood function for a given phenotype **y**, when evaluated at a point 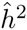, restated as a function of 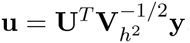. Fortunately, in several common scenarios, it is possible to give a closed-form expression of this derivative, whose computation is linear-time in the number of individuals.

We describe here the case where **K** was estimated from a standardized genotype matrix; and where **X** = **1_n_**, i.e. there are no covariates but there is an intercept. This is largely the most common use case. The full derivation for this case, as well as its extension to the case in which **X** additionally includes principal components of **K**, is given in Appendix A.4. This is a generalization of the work in [41, 42].

Let *d*_1_,…, *dn* be the eigenvalues of the kinship matrix in decreasing order, **K**, and recall that **U** is a matrix whose columns are **K**’s eigenvectors. For a true heritability value *h*^2^, a possible point estimate value *H*^2^, and for *i* = 1,…, *n* − 1, define

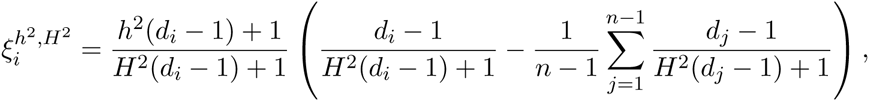

and also define 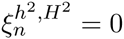.

A necessary condition for 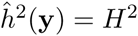, when 0 < *H*^2^ < 1, is having 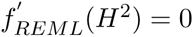 which translates to

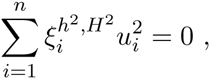

where 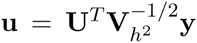 (a proof is given in A.4). Similarly, when 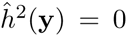 or 1, the respective conditions are having 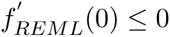 or 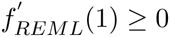, which translate to

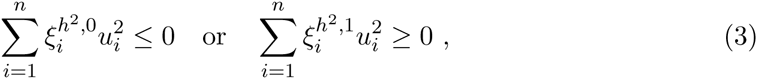

respectively. This is in line with the result of [41]. In practice, we only wish to bound the maximum within an interval. A sufficient condition for a local maximum to be in an interval (*c*_1_, *c*_2_) is having

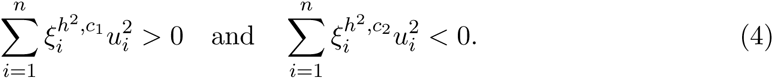

#### Step 3: Density estimation

To estimate the distribution of 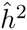, we draw 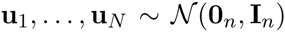. We then estimate the probability of 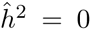 or 1 as the proportion of **u** values for which the respective Equations (3) hold. If the estimation for a certain **u***_i_* is neither 0 or 1, it means the derivative at 0 is positive and the derivative at 1 is negative. Therefore, we can perform a binary search to find a smaller interval (*c*_1_, *c*_2_) in which Equation (4) holds. This can be done until a small enough interval is found, at a specified resolution. In practice, we simply evaluate all intervals in a specified grid (e.g., (0, 0.01), (0.01, 0.02),…, (0.99, 1)).

A useful feature of this approach is that, in order to estimate the probability of a boundary estimate (e.g., 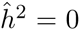) or an interval estimate (e.g., 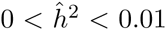), we need not find the maximum for each **u***_i_*, instead only checking if Equations (3), (4) hold.

### Confidence intervals for *h*^2^

We wish to build confidence intervals with a coverage probability of 1 − *α* (e.g., 95%). The distribution of 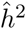 depends solely on *h*^2^, so we may assume without loss of generality that 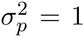 and *β* = 0*_p_* (see Appendix A.3). Our approach is based on the duality between hypothesis testing and confidence intervals. For a fixed value *h*^2^, an *acceptance region* 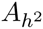 is defined as the subset of values 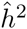 for which a test does not reject the null hypothesis that the phenotype is drawn from 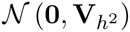. The probability of the event 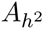 under 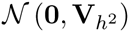 should be ≥ 1 − *α*. This region may be indirectly derived from an actual test (e.g., a generalized likelihood ratio test) or constructed explicitly. The corresponding confidence interval, 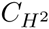, for an estimate 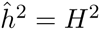 comprises of the set of parameter values for which the estimated values are not rejected, namely:

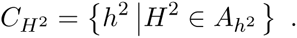

Since the distribution of 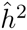 is bounded and generally asymmetric, the choice of 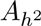 is not unique. It remains to determine 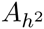 for every *h*^2^. We give here a general description of the construction; in Appendix A.5, we give a full description of the method, along with proofs.

Let _C_*_β_*(*h*^2^) be the *β*-th quantile function of 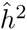, when the true heritability is *h*^2^. Since the distribution is of a mixed type with discontinuity points, it may be the case that 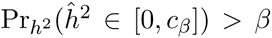. Specifically, the probability of the interval [*c_α_*_/2_(*h*^2^), c_1−_*_α_*_/2_(*h*^2^)] might be larger than (1 − *α*/2) − *α*/2 = 1 − *α*. We therefore divide our construction into distinct cases.

If there is a range of values *h*^2^ ∈ [*s, t*] for which 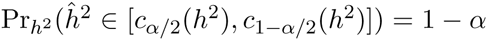, we set

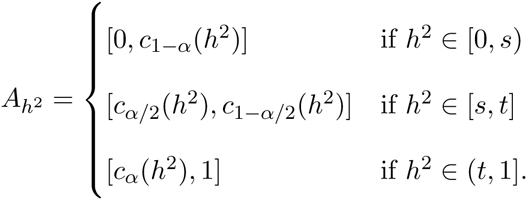

If no such range exists, but for every *h*^2^, either [0, *c*_1−_*_α_*(*h*^2^)] or [*c_α_*(*h*^2^), 1] have a probability of 1 − *α*, then set

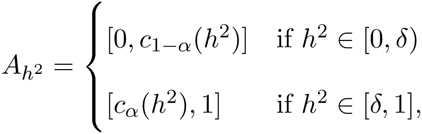

where *δ* is a point chosen so that the regions will have the required 1 − *α* probability. Finally, if there exists a value of *h*^2^ for which neither [0, *c*_1−_*_α_*(*h*^2^)] or [*c_α_*(*h*^2^), 1] have a probability of 1 − *α*, we employ randomized confidence intervals to achieve the required accuracy (see Appendix A.5).

### Benchmarks

We compared the computational costs of performing a parametric bootstrap procedure to estimate the heritability estimator distribution, using ALBI, the aforementioned GCTA [35] and pylmm [43], a fast and lightweight LMM solver for use in GWAS.

We generated the estimator distribution for *h*^2^ = 0.5 with these three methods. For pylmm and GCTA, we generated 100 or 1000 random phenotypes, and estimated the heritability for each phenotype using the respective programs’ estimation methods. Using the default flags, GCTA did not converge for ~7% of the samples. For ALBI, we estimated the distributions at a precision of 0.01 and 0.001, using 100 or 1000 random samples.

Timing for all methods did not include calculation of eigenvalues and eigenvectors. Running times are reported for the GTEx, LURIC and NFBC datasets. Runs were aborted after 8 hours. All programs were run on a 2.2GhZ, 64-bit Linux workstation with 64GB RAM.

### Datasets

#### The GTEx dataset

The Genotype-Tissue Expression (GTEx) [32] study is a sample and data resource designed to study the relationship among genetic variation, gene expression, and other molecular phenotypes in multiple human tissues. It provides a collection of multiple different tissues per donor, along with their genotypes. We use whole-genome data, collected by the Illumina HumanOmni5M-Quad BeadChip. Prior to quality control (QC), the data consist of 191 sampled individuals and 4,276,680 SNPs. We apply the recommended QC [32], after which 185 individuals and 3,575,877 SNPs remain. For heritability estimation, we use whole-blood gene expression profiles for 30,116 genes.

#### The LURIC dataset

The LUdwigshafen RIsk and Cardiovascular Health (LURIC) [33] study is a project contributing to the identification and assessment of environmental and genetic factors for cardiovascular diseases. The study consists of patients hospitalized for coronary angiography between 1997 and 2000 at a tertiary care center in Southwestern Germany. Quality control steps undertaken here include removing samples with < 95% call rate, SNPs with < 98% call rate, minor allele frequency of < 1%, or Hardy–Weinberg equilibrium test p-value of < 10^−4^. In addition, individuals for which the reported sex did not match the genotype-inferred sex, as well as individuals whose genotypes were manually observed to be outliers in an MDS plot, were removed. Moreover, from each pair of individuals with relatedness of more than 0.1875, only one was reserved. From these, only individuals for which lipid data were collected were used. This process resulted in 867 sampled individuals and 687, 262 SNPs. For heritability estimation, we use 102 lipid profiles.

#### The NFBC dataset

We analyze 5,236 individuals from the Northern Finland Birth Cohort (NFBC) data set, which consists of genotypes at 331,476 genotyped SNPs and 10 phenotypes [36]. The 10 phenotypes are triglycerides (TG), high-density lipoproteins (HDL), low-density lipoproteins (LDL), glucose (GLU), insulin (INS), body mass index (BMI), C-reactive protein (CRP) as a measure of inflammation, systolic blood pressure (SBP), diastolic blood pressure (DBP), and height.

### Discussion

We have presented ALBI, an efficient method for computing the distribution of the REML estimator of heritability and for constructing accurate confidence intervals. We showed that ALBI is significantly faster than standard parametric bootstrap approaches in computing the true estimator distribution, which otherwise require the explicit construction of random phenotypes and full estimation of heritability for each phenotype. In addition, unlike current methods, ALBI allows the computation of the probability of heritability estimates lying inside a specified interval (and particularly at the boundaries), without the need to estimate the entire distribution.

One of the main limitations of the methods currently used for heritability estimation is that the construction of CIs or standard errors is based on approximations that, as we have shown here, often do not hold in practice, resulting in unreliable CIs. In contrast, the CIs built by ALBI are accurate by construction, and ALBI can be used as an add-on to any of the current methods (e.g., GCTA [35], GEMMA [10]).

In addition, ALBI may be used as a practical approach to investigate the effect of various kinship matrix operations on the usefulness of the heritability estimator. For example, failing to exclude individuals with high relatedness introduces near-zero eigenvalues to the kinship matrix, whose effect on the estimator distribution can be tested. Similarly, it is possible to test if the common practice of adding the first PCs as fixed effects in the mixed model improves the quality of the estimator. Moreover, while we theoretically expect larger sample sizes to produce smaller CIs, the relationship between sample size and the shape of the spectrum of the kinship matrix is intricate, and must be studied and validated for each dataset individually, which may be done with ALBI.

We focused in this work on the estimation of heritability in the bounded interval [0,1], which is its natural domain. It is well known that the heritability estimator in this range is biased due to the bounded parameter space (see Figure S1). Some software packages for the estimation of heritability (e.g., GCTA) allow performing the optimization of the maximum likelihood or REML in an unbounded region. The rationale there being that even though a negative heritability is not meaningful, unbounded REML estimates are unbiased [38], and confidence intervals can be used to test whether the heritability is greater than a certain value, which is often the question of interest. However, this approach is problematic; a more natural solution for this task is using a method such as ALBI to compute reliable CIs for the bounded estimates. Furthermore, the domain is not well defined when the resulting kinship matrix is not positive definite.

A promising direction for future research is improving the efficiency of ALBI further. Utilizing efficient interval searches and hashing schemes is expected to improve the time complexity. Additionally, we note that at the core of the approximation is the estimation of the cumulative distribution function (cdf) of a generalized chi-square random variable (for boundary probabilities) or the joint distribution of two such variables (for non-boundary probabilities). Various approximations for these cdfs are available in the context of quadratic forms of normal variables [44], but to our knowledge, none apply to the generalized setting we have presented here.

The method proposed here can be extended in several directions. As the distribution of the heritability estimator is generally asymmetric and of mixed type, there are several ways to define the acceptance regions that are used to determine the CIs. Each such choice comes with its advantages and disadvantages, and it is possible that different choices may be more suitable in different settings. Additionally, we focused here on CIs for 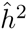 only. In practice, researchers may be interested in joint confidence regions for additional parameters, such as one or more of the fixed effects.

## Acknowledgements

The authors would like to thank Noah Zaitlen, Eli Levy-Karin and Yael Baran. This study was supported in part by a fellowship from the Edmond J. Safra Center for Bioinformatics at Tel-Aviv University to RS and SK. EH and SR are faculty fellows of the Edmond J. Safra Center for Bioinformatics at Tel Aviv University. RS and EH are partially supported by the Israeli Science Foundation grant 1425/13. SK and SR are partially supported by the Israeli Science Foundation grant 1487/12. RS is supported by the Colton Family Foundation. EH, SK, and RS are also partially supported by National Science Foundation grant III-1217615. EH, RS and EE are supported by National Science Foundation grant 1331176 and United States–Israel Binational Science Foundation Grant 2012304. The study was also funded by a European Union 7^th^ Framework Programme grant number 201668 for the AtheroRemo Project. EE is supported by National Science Foundation grants 1065276, 1302448, and 1320589 and National Institutes of Health grants R01-MH101782 and R01-ES022282. The data used for the analyses described in this manuscript were obtained from dbGaP accession number phs000424.v4.p1 on 4/4/2015.

## A Appendix

### A.1 Estimation of the kinship matrix

We follow the procedure described in the GCTA software package [1]. Let **Z** be a standardized genotype matrix with the *i, j*-th element 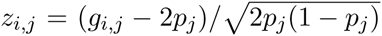, where *g_i,j_* is the number of copies of the reference allele for the *j*-th SNP in the *i*-th sample, and *p_j_* is the frequency of the reference allele, estimated from all non-missing SNPs. The *i*_1_, *i*_2_-th element of the kinship matrix is:

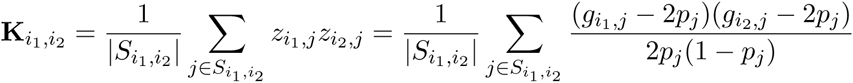

where 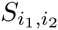 is the set of autosomal SNPs whose values are non-missing for both individuals *i*_1_ and *i*_2_. When there are no missing values, it follows that 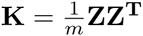.

### A.2 Estimating heritability, ML & REML

For the sake of completeness, we present the standard derivation of ML and REML estimation [2–4].

#### A.2.1 Maximum likelihood estimation

Define 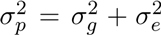 and 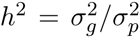. Up to additive and multiplicative constants, the log-likelihood function is

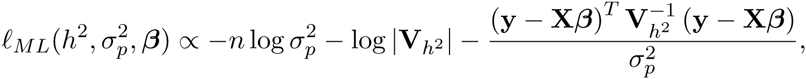

where 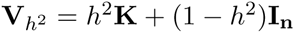. Therefore, 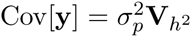. For a fixed *h*^2^, the values of 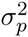 and *β* that maximize ℓ*_ML_* can be derived analytically:

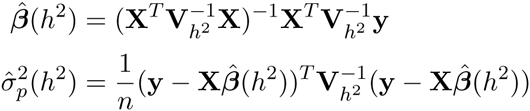

The problem of maximizing 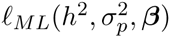 therefore reduces to maximizing the profile log-likelihood function 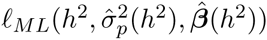 over *h*^2^, which is, up to constants,

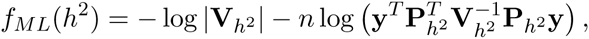

where 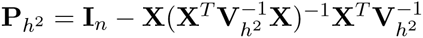.

#### A.2.2 Restricted maximum likelihood estimation

REML (Restricted Maximum Likelihood) was introduced to take into account the loss in degrees of freedom due to estimation of fixed effects [3, 5]. REML consists of maximizing the likelihood function that is associated with *n − p* linearly independent contrasts. The restricted log-likelihood function is, up to constants:

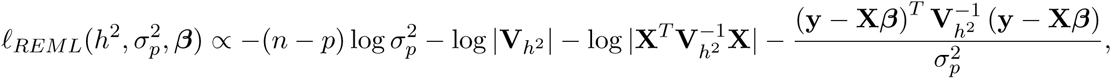

where now

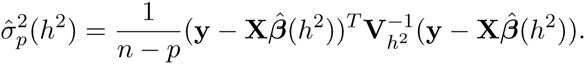

A similar analysis gives a profile function of:

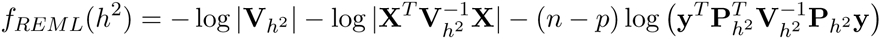

### A.3 Invariance of heritability estimates

We prove an invariance property of the likelihood and restricted likelihood functions, which enables us to focus solely on the estimator of *h*^2^ and disregard 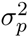 and *β*. For a fixed value of *h*^2^, the estimation problem reduces to standard regression with a covariance matrix that is known up to a scaling factor, 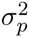. This leads to the generalized least squares solution for 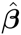, as it is independent of scale. It can be easily shown that, for 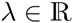 and 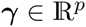:

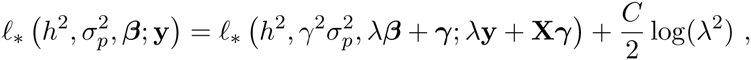

where *ℓ_*_* may be either *ℓ_ML_* or *ℓ_REML_*, and the constant *C* is *n* or *n − p*, respectively. Therefore, for 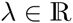 and 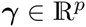:

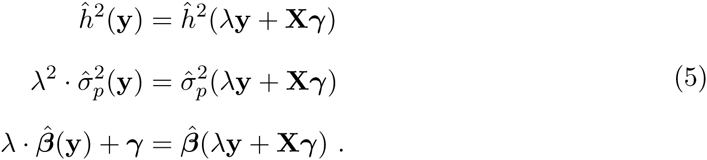

If **y** is drawn from the distribution of Equation (2) with parameters (*h*^2^, 1, **0_p_**), then 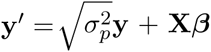 follows the distribution with parameters 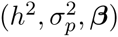. Therefore, from (5), the joint distribution of 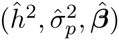, for any fixed 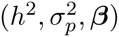, is determined exactly by the joint distribution of these estimators for (*h*^2^, 1, **0***_p_*). An important conclusion is that the distribution of 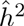 depends only on *h*^2^. We may therefore limit our study to the 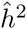 estimator alone, in the special case of fixed 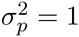 and *β* = **0***_p_*.

### A.4 The distribution of 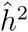

In [6], a necessary condition for estimating 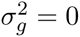 is derived, as the condition of the likelihood function having a non-increasing derivative at 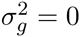. In addition, they approximate the probability of this event. We extend their work to derive a necessary condition for the ML/REML estimator evaluating to a general 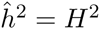. We then use this result to estimate the distribution of 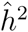.

#### A.4.1 Condition for estimating 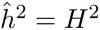, when 0 < *H*^2^ < 1

Let **K = UDU^T^** be the eigen-decomposition of **K**, where **D** is a diagonal matrix with the eigenvalues *d*_1_,…,*d*_n_, and **U** is an orthonormal matrix with the eigenvectors as its columns (If *d_i_* ≤ 0 (due to numeric errors), we round *d_i_* = 10^−10^ to make sure **K** is positive definite). Let

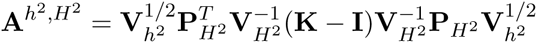

and

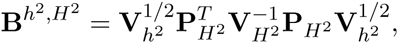

where 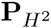 is defined in Section A.2.1. Let *w*_1_,…, *w_n_* be the eigenvalues of **KP_0_**, where 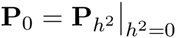. Let *p_i_* be an indicator variable determining if the *i*-th eigenvalue of **P**_0_ is 0. Define

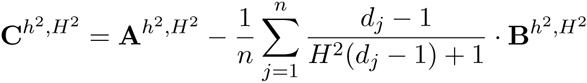

and

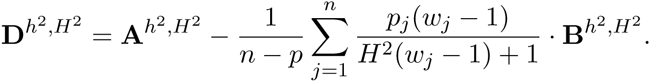

In addition, let 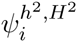 and 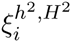 be the eigenvalues of 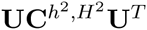 and 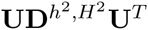, respectively, for *i* = 1,…, *n*. Finally, let 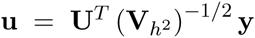. Then, the following holds:

##### Proposition 1.

*For* 0 < *H*^2^ < *1, a necessary condition for* 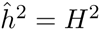 *for* **y** is:

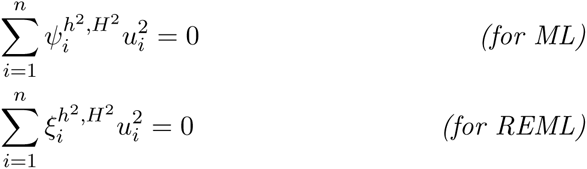

###### Proof.

A necessary condition for 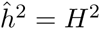 (when 0 < *H*^2^ < 1) being a global maximum for *f*_*_(*h*^2^) (where *f*_*_ is either *f_ML_* or *f_REML_*) is 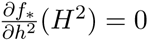. We show that this is equivalent to the equations above.

Utilizing Equations (7, 17) from [6], and applying the chain rule to convert these equations to be a function of *h*^2^ instead of 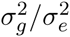, we get:

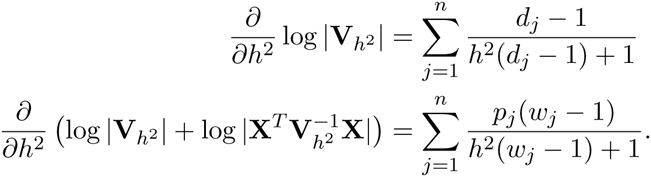

It is straightforward to verify that

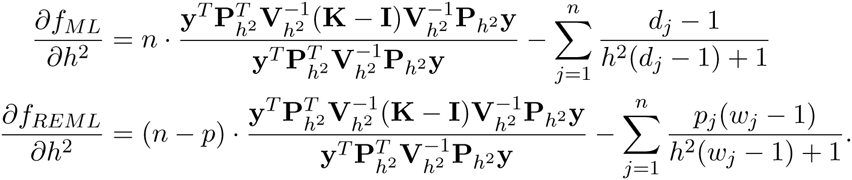

Let 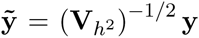. Then, by equating the derivatives to 0, it follows that a necessary condition for achieving a local maximum at *H*^2^ is

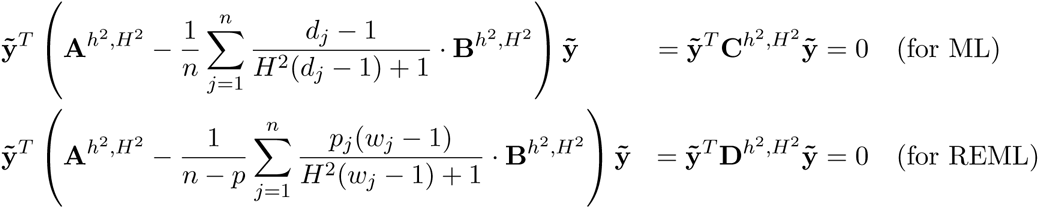

The eigenvalues of 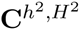 and 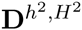 are denoted by 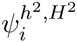 and 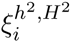, respectively (*i* = 1,…, *n*). Let 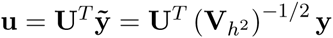. Then the conditions above are equivalent
to:

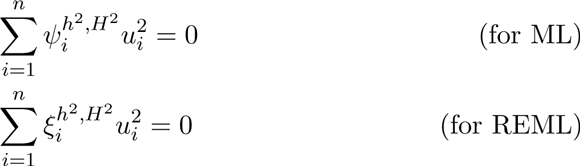

#### A.4.2 Condition for estimating 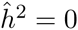 or 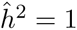

The following proposition shows that at the boundaries, the equality in the condition from Proposition 1 is replaced with an inequality.

##### Proposition 2.

*A necessary condition for* 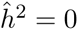 *is*

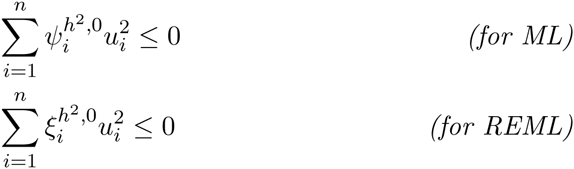

*Similarly, a necessary condition for* 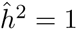 *is*

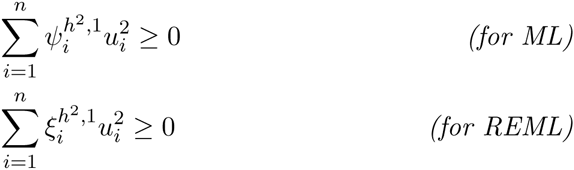

###### Proof.

For the case 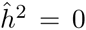, the requirement 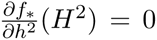 is replaced with 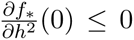, leading to the conditions above. Similarly, for the case 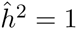, the requirement is 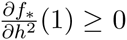.

#### A.4.3 Condition for 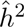 falling in a given interval

For 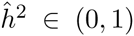, we will be estimating the distribution of 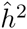 at a specified resolution. Therefore, we wish to know whether the estimate 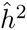 falls inside an interval (*c*_1_, *c*_2_). Using the derivative, it is simple to give a sufficient condition for an interval to contain a local maximum.

##### Proposition 3.

*A sufficient condition for the existence of a local maximum in* (*c*_1_, *c*_2_) *is*

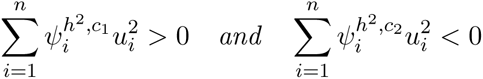

*for ML estimation, and*

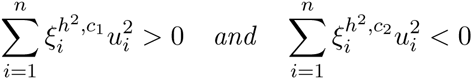

*for REML estimation.*

###### Proof.

If 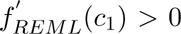 and 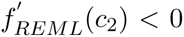, then there exists at least one 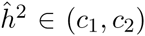 with 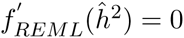 and 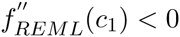. Therefore, a local maximum of 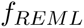 exists in (*c*_1_, *c*_2_). The case for ML is identical.

#### A.4.4 Linear-time simple formula for a class of X-s

The derivations above hold for a general covariate matrix **X**. Here, we limit ourselves to **X**-s whose columns are eigenvectors of **K**. Let **p** = (*p*_1_,…, *p_n_*) be an indicator vector specifying if the *i*-th eigenvector of **K** is not a column of **X**. This is a generalization of three common special cases:

1. **X** = 0. Namely, there are no fixed effects. The requirement about **X** holds trivially. Here, **p = 1_n_**.
2. **Intercept**. In many cases, **X = 1***_n_*, an intercept vector (e.g., the default in GCTA). Since each column of **Z** is standardized, **Z***^T^* **1***_n_* = **0***_m_*, so 1*_n_* is an eigenvector of **K**, corresponding to an eigenvalue of 0. Since **K** is non-negative definite, this is the lowest eigenvalue, i.e. *d_n_* = 0. In practice, this hold only approximately due to numerical errors in computing the eigenvalues, but the approximation is good enough for this analysis to be useful. Here, 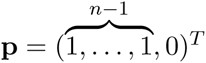.
3. **First PCs**. In addition to an intercept term, a common practice is adding to **X** the *q* largest principal components, corresponding to the first *q* eigenvectors of **K**. Here,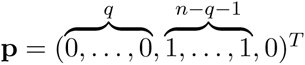

In these cases, it is possible to present a simpler formula.

##### Proposition 4.

*If the columns of* **X** *are eigenvectors of* **K***, then*

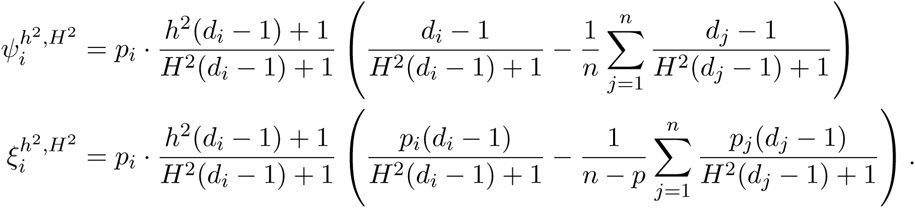

###### Proof.

Recall that **K = UDU^T^** is the eigen-decomposition of **K**, with the eigenvalues *d*_1_,…, *d_n_*. It follows that 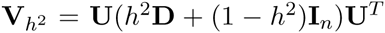. 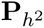 is the projection matrix to the subspace orthogonal to the subspace spanned by the columns of **X**. Therefore, 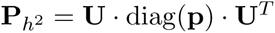 (and thus this definition of *p_i_* coincides with the one above). Recall also that

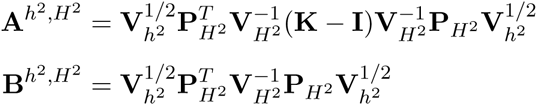

and denote their respective eigenvalues *a_i_* and *b_i_* Since all the matrices in these expressions are diagonalizeable by **U**,

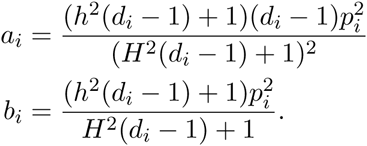

It follows that

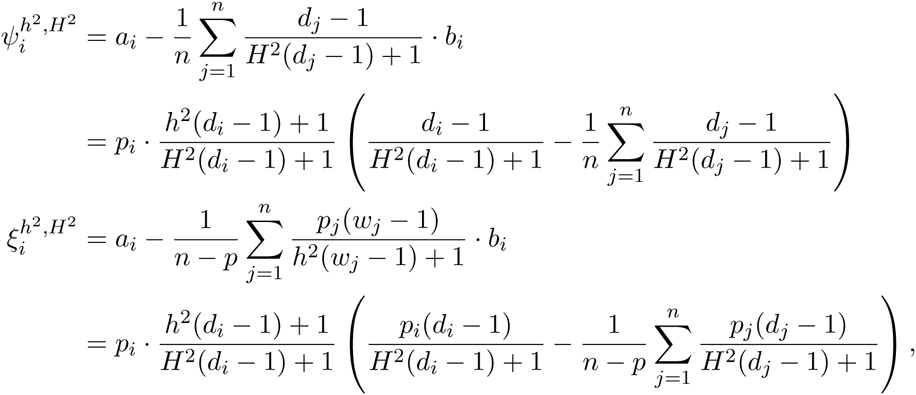

where we used 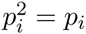 and that the eigenvalues of **KP_0_** are *w_j_ = d_j_p_j_*.

Once the eigenvalues of **K** are given, the calculation of 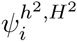 and 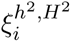 is performed in O(*n*) complexity. This allows for a simple closed-form formula and fast evaluation of the condition for maximality, even for large datasets.

### A.5 Construction of confidence intervals

We wish to build a set of confidence intervals with a coverage probability of 1 − *α* (e.g., 95%). The distribution of 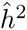 depends solely on *h*^2^, so we may assume without loss of generality that 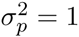, ***β* = 0***_p_* (see A.3). Our approach to CI construction is based on the duality between hypothesis testing and CIs (for a comprehensive treatment of this subject, see [7]).

Let 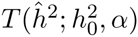 be any test for the null hypothesis 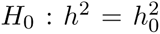, which has size *α* (i.e., whose type-I error rate is no more than *α*), and which employs 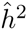 as its test statistic. Let 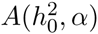 be the acceptance region of this test, i.e., 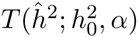 does not reject *H*_0_ iff 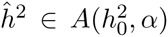. Since the test is of size *α*, we know that, under the null, 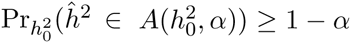. The CI (or, more generally, confidence set) 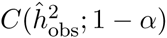 that is dual to the test 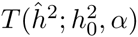 simply comprises of all parameter values *h*^2^ not rejected by *T* when observing the estimate value 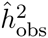, i.e., 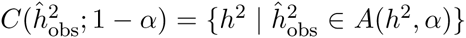. From this definition, we have that

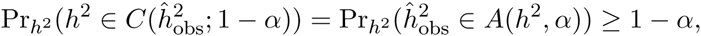

namely, *C* is a CI with coverage probability at least 1 − *α*. Moreover, if the acceptance region has null probability 1 − *α* exactly (i.e., its associated test is not conservative), then the CI achieves coverage 1 − *α* accurately (otherwise, the CI is also conservative). In the following, we simplify the notation for the CI to 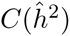, leaving the dependence on *α* implicit. We describe a method to construct accurate acceptance regions, and therefore confidence intervals, with coverage probability of 1 − *α*.

#### A.5.1 Preliminaries and Assumptions

Define:

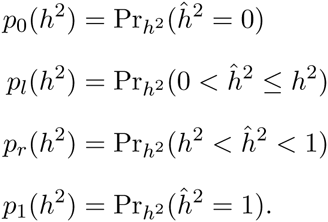

To prove the accuracy of the CIs, we assume several natural properties of the distribution of the estimators:

##### Assumption 1.

*The boundary probabilities p*_0_(*h*^2^) *and p*_1_(*h*^2^) *are monotone decreasing and increasing functions of h*^2^, *respectively.*

Figure 4 in the main text indicates that Assumption 1 holds in practice.

##### Assumption 2.

*The required confidence level is larger than the maximum of the boundary probabilities; namely,* 1 − *α* > max(*p*_0_(0), *p*_1_(1)).

To see why Assumption 2 is reasonable, note that the largest boundary probabilities *p*_0_(0), *p*_1_(1) are expected to be ~0.5 asymptotically [8]. Indeed, similar values of ~0.5 − 0.6 are also seen in practice (see Figure 4 in the main text). Meanwhile, 1 − *α* is the required confidence level, with 0.95 being a typical value.

For every *h*^2^ ∈ [0, 1], the open interval (0, 1) is mapped bijectively to (*p*_0_(*h*^2^), 1 − *p*_1_(*h*^2^)) by the CDF 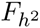. The quantile function c*_β_* of 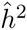 is:

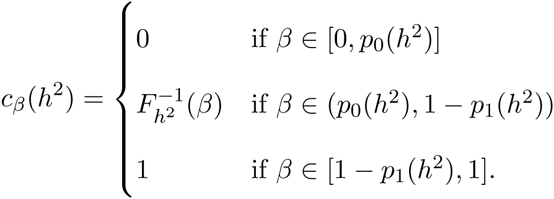

Note that since estimator distributions are discontinuous at the boundaries, it may be the case that *c_β_*(*h*^2^) does not obey 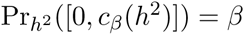.

##### Assumption 3.

*For every β,* c*_β_*(*h*^2^) *is non-decreasing in h*^2^.

It follows that

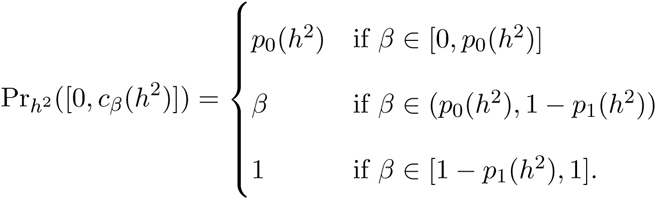

##### Assumption 4.

For *all h*^2^ ∈ [0,1], *h*^2^ ≤ *c*_1−_*_α_*(*h*^2^) and *h*^2^ ≥ *c_α_*(*h*^2^).

While this may not hold if 1 − *α* < *p*_0_(*h*^2^), it holds in practice for all common values of 1 − *α* (e.g., 95%). Note that since by definition, *c*_1−_*_α_*(*h*^2^) ≤ *c*_1−_*_α_*_/2_(*h*^2^) and *c_α_*(*h*^2^) ≥ *c_α_*_/2_(*h*^2^), it follows that *h*^2^ ≤ *c*_1−_*_α_*_/2_(*h*^2^) and *h*^2^ ≥ *c_α_*_/2_(*h*^2^).

#### A.5.2 Acceptance regions

A natural requirement of confidence intervals is that their lower and upper bounds be monotone increasing functions of 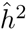. Equivalently, we require the lower and upper bounds of the acceptance regions to be monotone increasing functions of *h*^2^. Therefore, acceptance regions for increasing values of *h*^2^ are naturally divided into three types: Regions with lower bound 0; then, as the lower bound increases, possibly regions with both upper and lower bounds between 0 and 1; and finally, regions with an upper bound of 1. Specifically, we define the three types of regions to be:

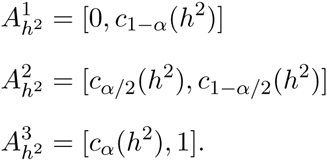

The three types are defined so that 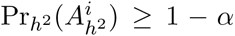. It is possible that equality is not obtained. For example, if *p*_1_(*h*^2^) > *α* at a point *h*^2^, then *c*_1−_*_α_*(*h*^2^) = 1, and then 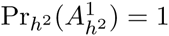.

#### A.5.3 Choosing regions

We now define how to choose a region 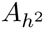 for each value of *h*^2^ from these three types. There are three requirements from this choice of regions that are desirable:

1. For the respective CIs to be accurate, 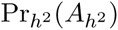 must be equal to 1 − *α*.
2. The true parameter value, *h*^2^, must be included in each region 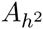.
3. As previously mentioned, the lower and upper bounds of the regions must remain monotone functions of *h*^2^.

By construction, moving from smaller values of *h*^2^ to larger ones, we consecutively use regions of type I, possibly of type II, and then type III. By definition, we choose a type I region for *h*^2^ = 0 and type III for *h*^2^ = 1. There are three scenarios we might encounter:

**1. Type II is used**. Suppose there is a range of values *h*^2^ ∈ [*s, t*] for which 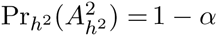. Then, we pick *s* and *t* as the transition points, i.e.,

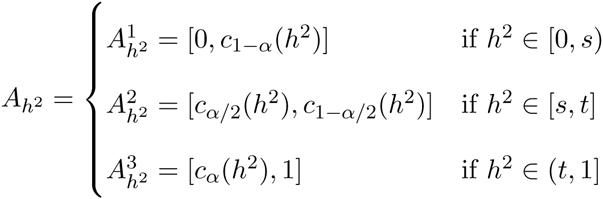

This choice of transitions maintains the monotonicity of the lower and upper bounds of the regions. To see this, note that within each of the parameter ranges, [0, *s*), (*s, t*) and (*t*, 1], monotonicity is obeyed via Assumption 3. At the transition points, monotonicity is maintained by definition of *c_β_*, with *c*_1−_*_α_*(*h*^2^) ≤ *c*_1−_*_α_*_/2_(*h*^2^) and *c_α_*_/2_(*h*^2^) ≤ *c_α_*(*h*^2^). Also, from Assumption 4, 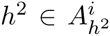 for *i* = 1, 2, 3. Therefore, 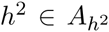 holds for all *h*^2^. The following proposition shows that the CI is accurate.

##### Proposition 5.

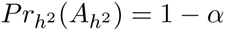 *for all h*^2^ (*i.e. the CI is accurate*).

###### Proof.

If *β* > *p*_0_(0), then since *p*_0_ is monotone decreasing (Assumption 1), *β* > *p*_0_(*h*^2^) for all *h*^2^. Therefore, for such values of *β*, 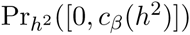 obtains only the values *β* (when *β* < 1 − *p*_1_(*h*^2^), or equivalently, when 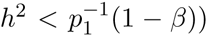 and 1 (when *β* ≥ 1 − *p*_1_(*h*^2^), or equivalently, when 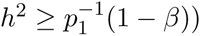.

Since 1 − *α* > *p*_0_(0) (Assumption 2), 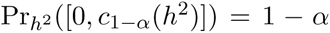 iff 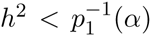 and 1 otherwise. Also, since 1 − *α*/2 > 1 − *α* > *p*_0_(0), similarly 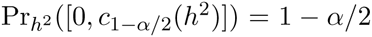 iff 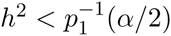 and 1 otherwise.

Since in the range *h*^2^ ∈ [*s, t*], 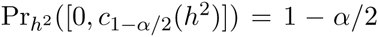, it follows from the above that 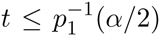. Since *p*_1_ is monotone increasing (Assumption 1), it follows that 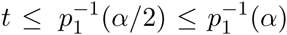. Therefore, in the range [0, *s*) ⊂ [0, *t*), the type I region 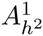 has the required probability, 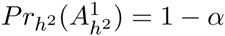.

A similar argument holds for the other direction. If 1 − *β* > *p*_1_ (1), and since *p*_1_ is monotone increasing, *β* < 1 − *p*_1_(*h*^2^) for all *h*^2^. Therefore, 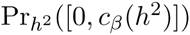 equals *p*_0_(*h*^2^) when 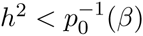 and *β* when 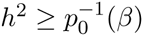.

Since in the range *h*^2^ ∈ [*s, t*], 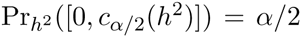, then 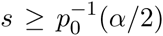. Since *p*_0_ is monotone decreasing, 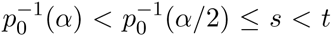. Therefore, in the range (*t*, 1], 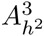 has the required probability, 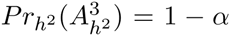. Consequentially, 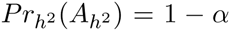 for all *h*^2^.

**2. Type II is not used, but types I and III achieve accuracy**. If there is no range where 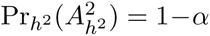, we turn to see if regions of either types I and III may have a probability of 1 − *α* for all *h*^2^ ∈ [0,1]. We have seen that 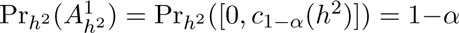 iff 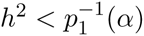 and 1 otherwise. We have also shown that 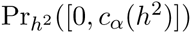 equals *p*_0_(*h*^2^) when 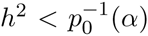 and *α* when 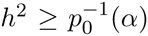. It follows that 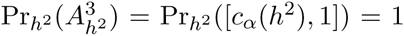 when 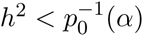 and 1 − *α* when 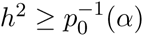.

Therefore, if 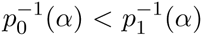, there does not exists a value *h*^2^ for which neither 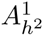 nor 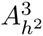 have the required probability of 1 − *α*. We pick their average 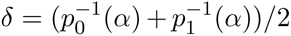 as the transition point, i.e.,

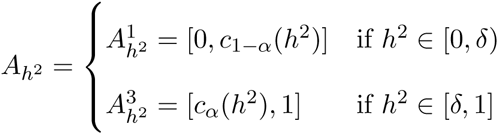

Consequentially, 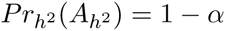 for all *h*^2^. Monotonicity of region bounds is trivial in this case, and again, from Assumption 4, 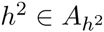 holds for all *h*^2^.

**3. Type II is not used, and types I and III do not achieve accuracy**. Finally, if there is no range where 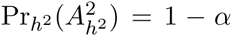, and 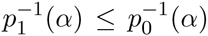, then in the range 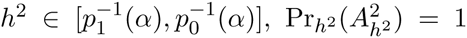 for all three region types. In this case, we use randomized confidence intervals [9] to achieve the exact required accuracy.

In this case, it is simpler to discuss the confidence intervals directly. Let

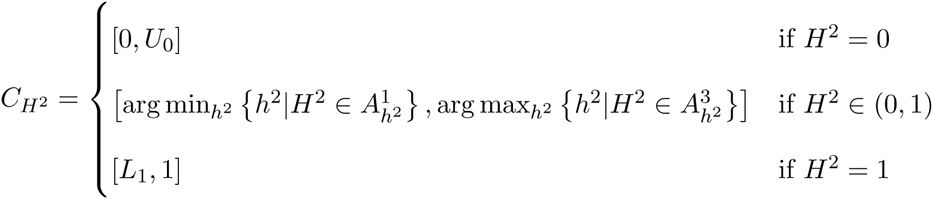

where *U*_0_ and *L*_1_ are random variables defining the upper bound of *C*_0_ and the lower bound of *C*_1_, respectively. Let 1 − *u*(*h*^2^) be the cdf of *U*_0_, and let *l*(*h*^2^) be the cdf of *L*_1_. It remains to choose *u*, *l* that will be appropriately monotone and will give the required coverage. The total coverage should obey:

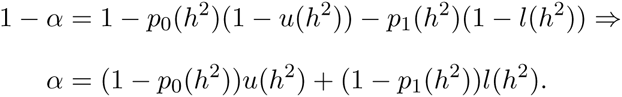

The requirement *U*_0_, 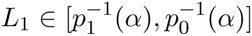 translates to:

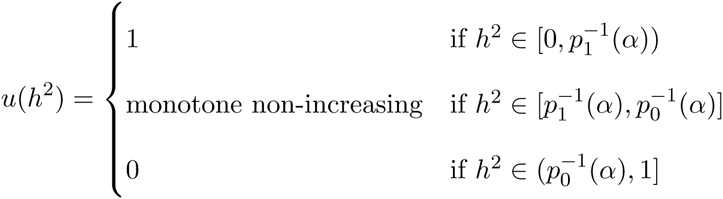

and

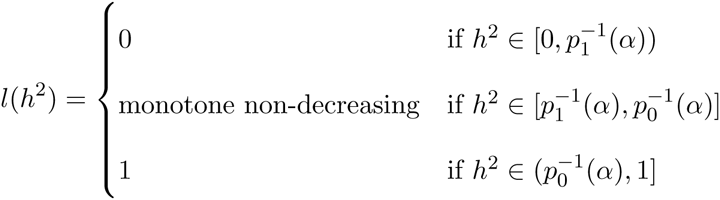

Inverting our viewpoint back to acceptance regions, the above is equivalent to choosing 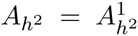 when 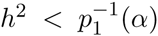 and 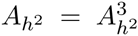 when 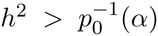. When 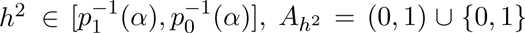, with the boundary points 0 and 1 included with probabilities *u*(*h*^2^) and *l*(*h*^2^) respectively.

When *l*(*h*^2^) = 1,

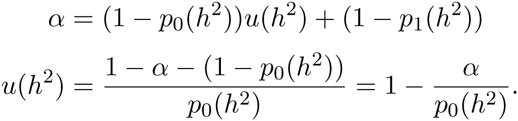

Similarly, when 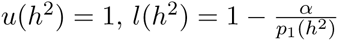.

The particular form we choose for *u* and *l* is

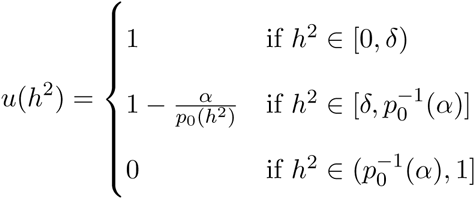

and

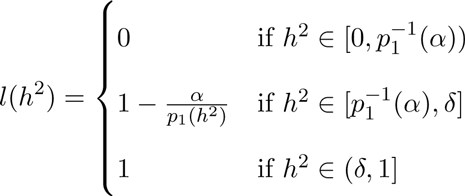

where 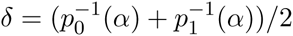, as before.

### A.6 Variance of estimators

The main method of calculating the variance of the estimator, applied by all widely used LMM methods, employs the Fisher information matrix, or a variant of which, possibly applying the delta method in addition [10]. The (expected) Fisher information matrix of an estimator 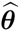 is the matrix 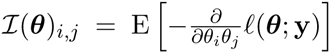. Asymptotically, under certain regularity conditions, 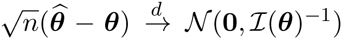 According to the delta method, the asymptotic distribution of a function *f*(***θ***) satisfies 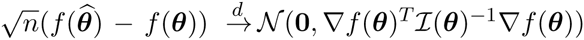.

GCTA uses the Average Information [11] (AI) matrix to calculate the variance of 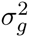 and 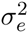. The AI matrix is an average between the expected and the observed Fisher information matrices. For the REML method, this is the matrix:

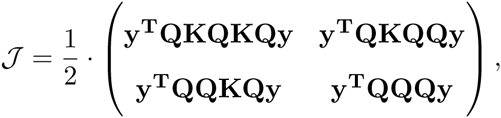

where **Q = Σ^−1^** − **Σ^−1^ Χ (X***^T^***Σ**^−^**^1^Χ***^T^*)^−^**^1^ X***^T^***Σ**^−^**^1^**, with 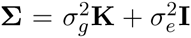. Then, the delta method is used to calculate the variance of 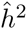:

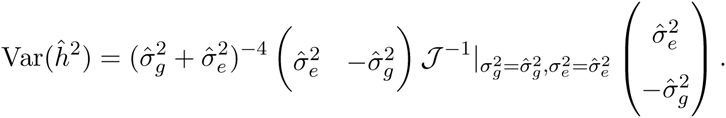

GEMMA instead applies the delta method on a closed-form formula of the ratio 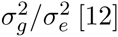.

## Supplementary Figures and Tables

### REML estimator bias

We evaluated the empirical bias of the REML estimator 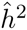 as a function of *h*^2^. It is evident that the REML estimator may be biased (Figure S1). The bias is especially evident for low and high true values of *h*^2^. This is also true for the estimator of 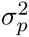 (not shown).

**Figure S1.**
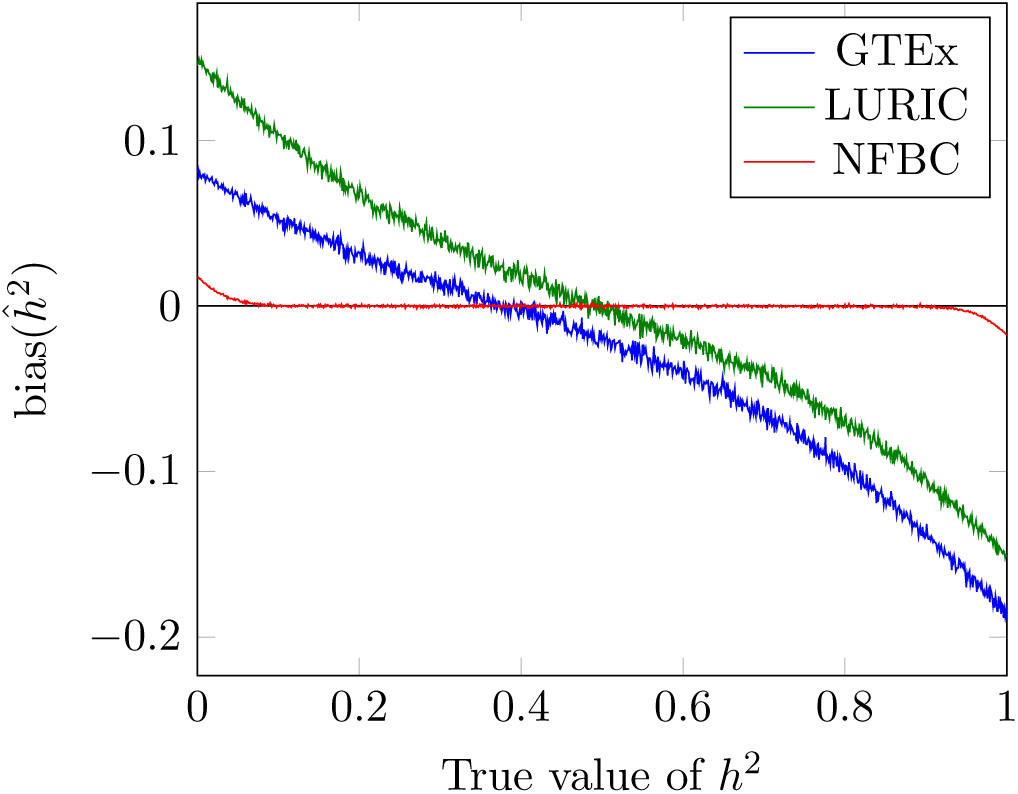
Bias of the heritability estimator. The bias of 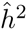 (defined as 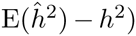, for all possible values of true underlying heritability *h*^2^, for the studied datasets. The GTEx and LURIC datasets create highly biased estimators, while for the NFBC the estimator is mostly unbiased.

